# Disease mortality in domesticated animals is predicted by host evolutionary relationships

**DOI:** 10.1101/438226

**Authors:** Maxwell J. Farrell, T. J. Davies

## Abstract

Infectious diseases of domesticated animals impact human well-being via food insecurity, loss of livelihoods, and human infections. While much research has focused on parasites that infect single host species, most parasites of domesticated mammals infect multiple species. The impact of multi-host parasites varies across hosts; some rarely result in death, whereas others are nearly always fatal. Despite their high ecological and societal costs, we currently lack theory for predicting the lethality of multi-host parasites. Here, using a global dataset of over 4000 case-fatality rates for 65 infectious diseases (caused by micro and macro-parasites) and 12 domesticated host species, we show that the average evolutionary distance from an infected host to other mammal host species is a strong predictor of disease-induced mortality. We find that as parasites infect species outside of their documented phy-1 logenetic host range, they are more likely to result in lethal infections, with the odds of death doubling for each additional 10 million years of evolutionary distance. Our results for domesticated animal diseases reveal patterns in the evolution of highly lethal parasites that are difficult to observe in the wild, and further suggest that the severity of infectious diseases may be predicted from evolutionary relationships among hosts.

## Introduction

Infectious diseases that cross species barriers are responsible for severe human health burdens (Hotez et al., 2014), and act as direct and synergistic drivers of species extinctions (Heard et al., 2013). Many of these diseases infect domesticated animals and impact human well-being via loss of food security, labour and livelihoods, costs of prevention and control programs, and increased human infection (Dehove et al., 2012). However, the severity of disease can vary dramatically among parasites. Ca-nine rabies alone results in approximately 59,000 human deaths and 8.6 billion USD in economic losses annually (Hampson et al., 2015). By contrast, other diseases rarely result in death. For example, bovine brucellosis largely impacts cattle by causing abortion, infertility and reduced growth, but disease induced mortality in adult cows is uncommon (McDermott et al., 2013).

Well established theory on single-host parasites predicts that the reduction in host fitness due to infection (termed “virulence”) should evolve to an optimal level determined by a trade-off with transmission (Cressler et al., 2016). For multi-host parasites, optimal virulence may be subject to additional trade-offs, with selection for high or low virulence depending on the ecologies and evolutionary histories of each susceptible host species (Woolhouse et al., 2001; Gandon, 2004; Rigaud et al., 2010). In the absence of trade-offs, a wider host breadth should provide a larger pool of susceptible individuals, increasing opportunities for transmission and the evolution of higher virulence (Barrett et al., 2009). However, adaptation to novel hosts may reduce a parasite’s ability to utilize resources of their co-evolved hosts (Ebert, 1998; Longdon et al., 2014), resulting in limited replication and decreased virulence (Antonovics et al., 2013). This trade-off is supported by comparative studies of plant RNA viruses and avian malaria parasites in which specialist parasites tend to be more virulent than generalists (Garamszegi, 2006; Agudelo-Romero and Elena, 2008). Yet generalist parasites remain highly virulent in some host species (Leggett et al., 2013).

Our ability to predict the outcome of a given host-parasite interaction is currently limited because the full suite of traits underlying virulence is either poorly estimated or unknown for the vast majority of host-parasite interactions. However, our understanding of evolutionary relationships is often much better, and host phylogeny can be used as a proxy for latent traits and evolutionary histories that have shaped contemporary host-parasite associations (Davies and Pedersen, 2008). For example, closely related hosts suffer similar impacts for some parasites of *Drosophila* (Longdon et al., 2015; Perlman and Jaenike, 2003), consistent with the prediction that parasite virulence should co-vary with host phylogeny. However, there have been few studies that develop and test theories of how host evolutionary relationships influence disease outcomes across multiple host-parasite combinations.

As parasites adapt to infect novel host species increasingly distant from their co-evolved hosts, they are expected to experience increased fitness costs (Antonovics et al., 2013), leading to the prediction of lowered virulence following greater phylogenetic jumps. This pattern, termed “non-host resistance” (Antonovics et al., 2013), may act in opposition to resistance evolved by hosts in response to infection, which is expected to decrease with evolutionary distance from a parasite’s co-evolved hosts and lead to phylogenetically distant hosts experiencing more intense disease (Antonovics et al., 2013). The relative strengths of these opposing relationships will likley influence the virulence of a given host-parasite interaction.

Infectious diseases of domestic species, many of which have severe economic impacts (Dehove et al., 2012), present a unique opportunity to explore the links between virulence, host specificity, and the evolutionary relationships among hosts. While virulence can take many forms, mortality is most widely reported. The World Organisation for Animal Health (OIE) publishes yearly reports documenting the numbers of cases and deaths caused by diseases of importance for international trade (for Animal Health (OIE), 2016), providing a remarkable dataset of disease-induced mortality for multiple parasites across different host species. We examine data from 4157 reports (in which no host culling was recorded) from 155 countries across 7 years, representing 202 unique host-parasite combinations with large variation in average mortality (Fig 1A).

**Figure 1:**
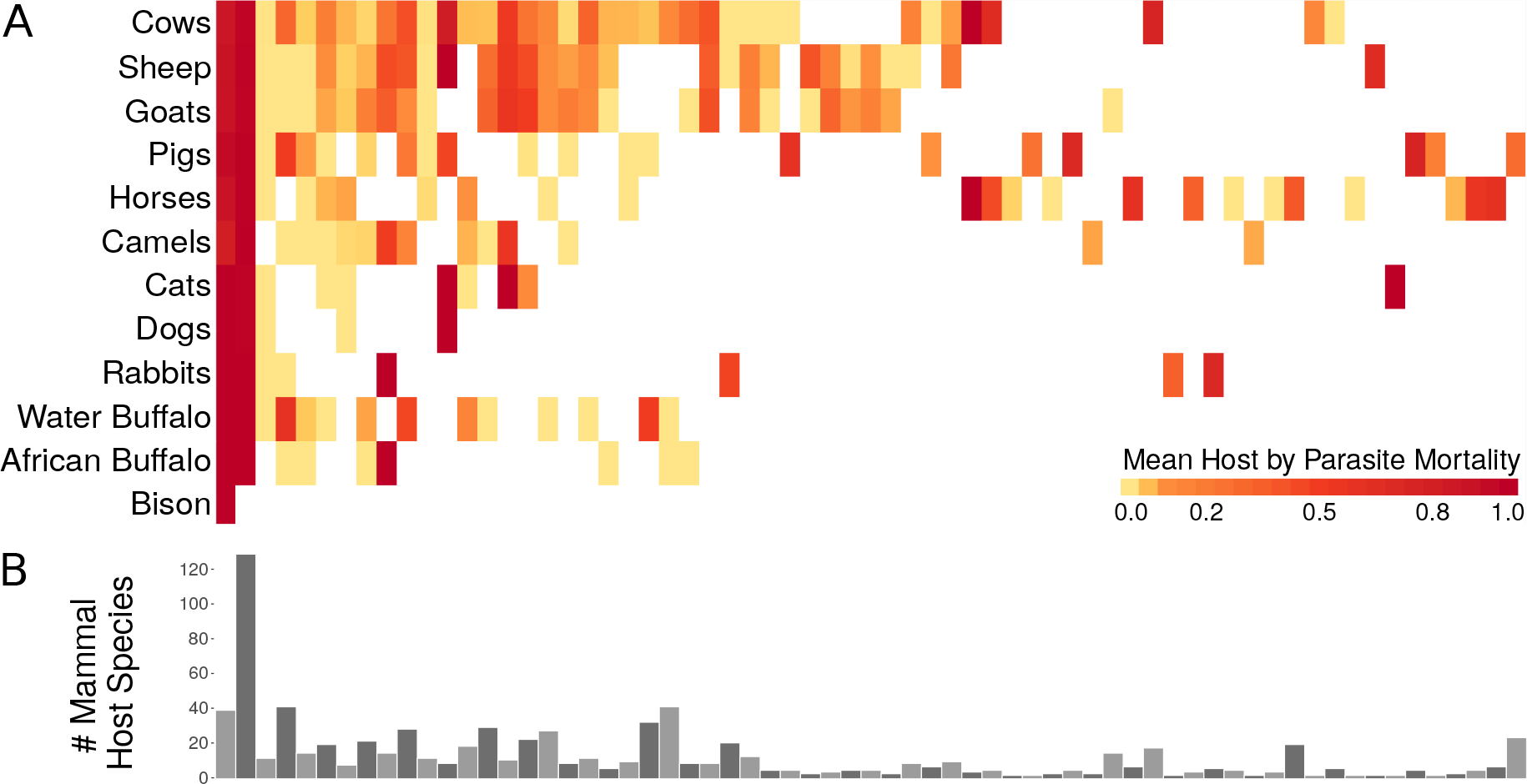
A) Heatmap of mean host by parasite mortality derived from the OIE World Animal Health yearly reports from 2005-2011 (full heatmap with disease common names included in Supplementary Fig 7). B) Barplot of the number of documented mammal host species per parasite derived from the Global Mammal Parasite 2.0 and EID2 databases. Order of parasites matches column order in B).

For each parasite, we identified the set of documented mammal host species from two recently published global host-parasite databases (Stephens et al., 2017; Wardeh et al., 2015), returning 788 unique host-parasite interactions (Fig 1B). For each host-parasite combination, we then calculated the mean phylogenetic distance from all documented host species to the infected species, which we refer to as “host evolutionary isolation” (Fig 2). This metric is analogous to measures of mean phylogenetic relatedness to a focal species, which have been used to analyze species invasions (Strauss et al., 2006), and predict disease pressure in plant communities (Parker et al., 2015). Since parasites typically infect closely related species (Antonovics et al., 2013; Davies and Pedersen, 2008), we assume the phylogenetic centroid of susceptible species indicates the likely position of ancestral hosts, and that the distance from a host to this centroid may provide a reasonable proxy for the relative extent of co-adaptation between parasite and host. We modelled the probability of death as a function of host evolutionary isolation and number of documented host species (host species richness) using a hierarchical Bayesian approach (The Stan Development Team, 2017) that allowed us to control for additional factors including the number of cases per report, and the effects of parasite, host, country, and year.

## Results

We find that disease-induced mortality is highest when infected hosts are evolutionarily distant from other documented hosts (Fig 3A, Fig 4, Table 2), with an increase of 10 million years of evolutionary isolation resulting in a doubling in the odds of host death (odds ratio 50% credible interval: 1.99 - 2.15). This predicts that a parasite infecting an Artiodactyl, which otherwise infects only Primate hosts, would have ∼ 4.8 times higher odds of host death than a parasite that otherwise infects hosts in the order Carnivora. This effect size is comparable in magnitude only to the number of cases, and much greater than the effect of all ecological and socioeconomic predictors in our model. The effect of host evolutionary isolation becomes stronger when single-host parasites are excluded (Table 3), indicating the results are not driven simply by differences between single and multi-host parasites.

We find some support for a positive relationship between mortality and host species richness (50% credible interval does not overlap zero), opposite to what would be predicted if there was a trade-off between parasite generalism and virulence (parasites with larger host richness causing lower mortality). However, there is large variability in the strength of this relationship, as is the case for all parasite-level predictors. Reports with high host mortality were associated with fewer infected individuals (Fig 3B, Fig 4), consistent with the upper extreme of the virulence-transmission trade-off. High mortality may naturally limit transmission, however human interventions to limit spread may also be strongest for deadlier outbreaks in domesticated animals.

**Figure 2:**
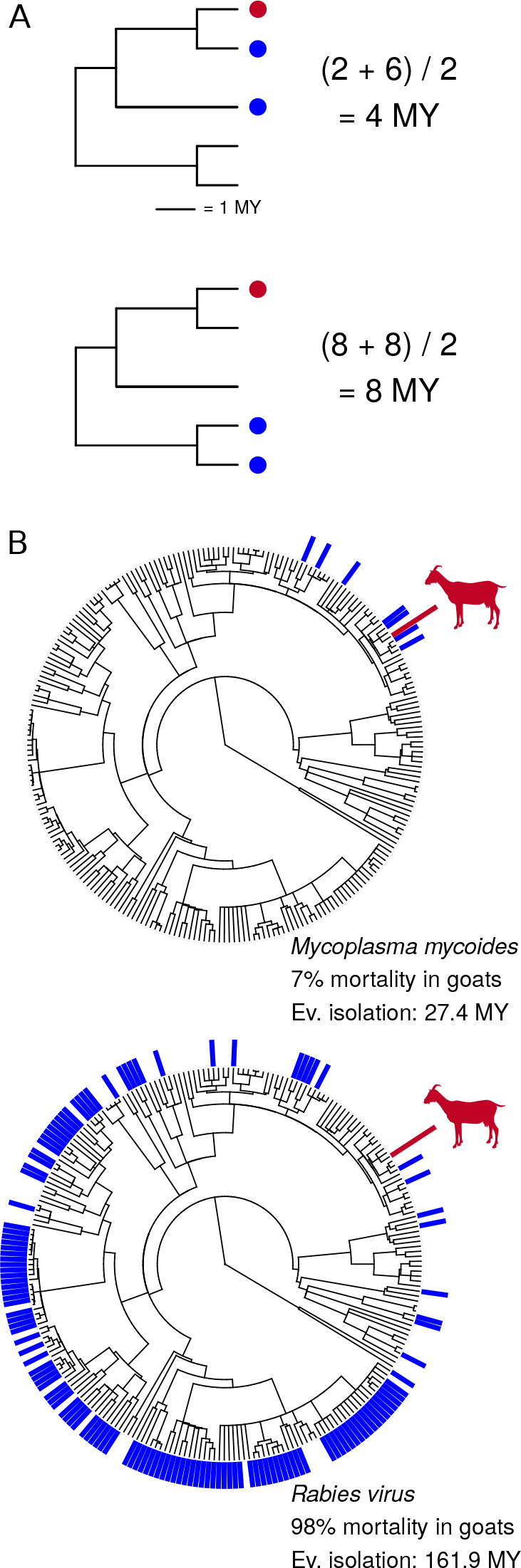
A) Example of how host evolutionary isolation is calculated. Red circles indicate the infected host, blue circles indicate documented hosts. Host evolutionary isolation is calculated as the mean phylogenetic distance from the infected host to all documented host species. B) Examples with *Mycoplasma mycoides* and *Rabies virus*. Documented hosts are indicated by blue bars on the host phylogeny, with host evolutionary isolation and average mortality calculated for goats (*Capra hircus*, shown in red).

Our model also revealed large variation in mortality among countries (Fig 3, Fig 5), indicating that effective disease management practices from one nation could be identified and introduced to other nations. Countries with large positive effects – higher mortality than otherwise predicted – may have lower capacities for detection and prevention of outbreaks. For example, top ranked Sri Lanka and Kyrgyzstan have struggled to develop legislation and infrastructure for addressing veterinary public health issues (Dissanayake et al., 2012), and have deteriorated veterinary and sanitation systems (Counotte et al., 2016). In contrast, nations with large negative country effects (the former Yugoslav Republic of Macedonia, China, and Iran) suffer considerable infectious disease burdens, but have made great improvements in surveillance, control, and eradication programs (Stojmanovski et al., 2014; Hotez et al., 2012; Wang et al., 2008). In addition, we found support for a negative relationship between mortality and GDP per capita (Fig 4), indicating that wealthier countries may allocate greater resources towards animal health and disease control efforts.

**Figure 3:**
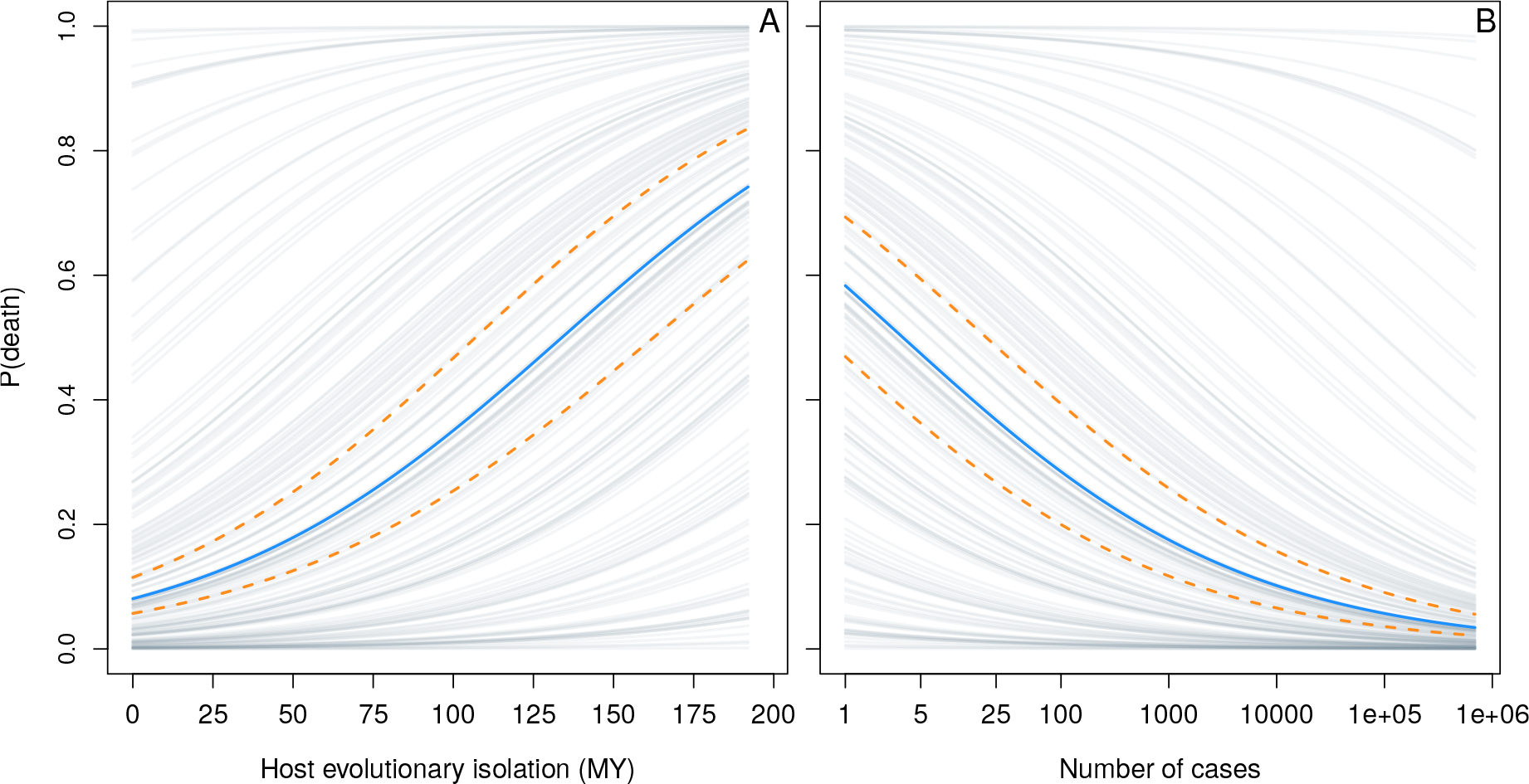
Posterior predictions of the probability of death as a function of A) host evolutionary isolation (in millions of years), and B) the number of cases. Solid blue lines represent the mean logistic curve, dashed yellow lines represent the upper and lower bounds of the 50% credible interval. Grey lines depict equivalent mean curves offset by the posterior mean effects for each country.

## Discussion

We find that as parasites infect domesticated species outside of their typical evolutionary host range, they have a higher probability of resulting in lethal infections. However, high mortality is also associated with fewer infected individuals. Our findings suggest that disease spillover into evolutionary 114 isolated hosts is marked by increased virulence, but potentially at the cost of decreased transmission. The high mortality observed in our data likely occurs through multiple pathways including the maladaptation of both host and parasite, and the decoupling of transmission from virulence. While it is difficult to determine the precise mechanisms leading to high mortality, we suggest that the evolutionary distances among infected and susceptible hosts can, to some extent, capture these multiple dimensions.

**Figure 4:**
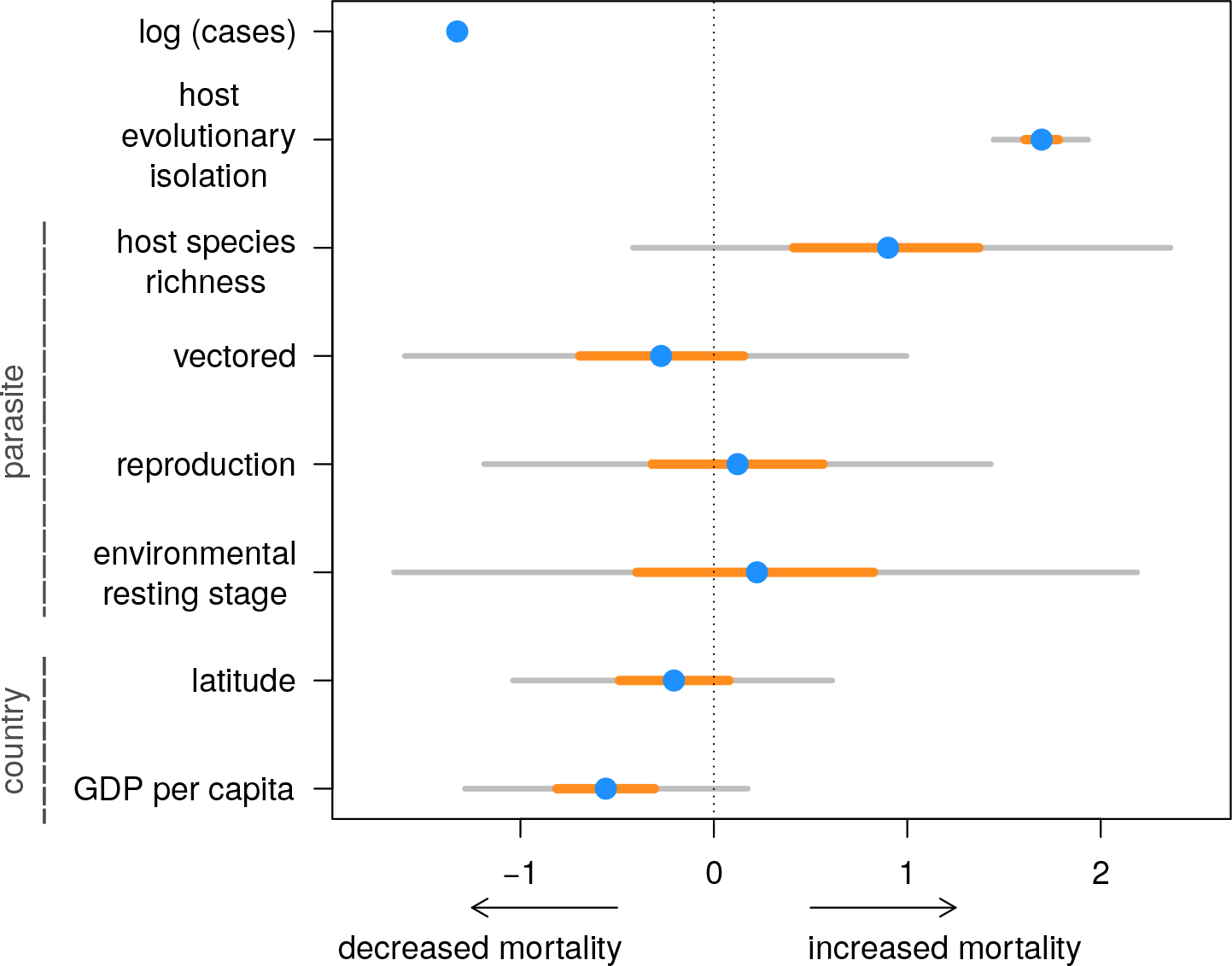
Estimated regression coefficients for continuous predictors. Blue circles represent posterior means, yellow horizontal lines represent 50% credible intervals, gray horizontal lines represent 95% credible intervals. Predictors at the parasite and country level are indicated.

For some host-parasite combinations, elevated mortality may be explained by a decoupling of virulence from transmission. Consistent with this hypothesis, many vertebrate arboviruses commonly use birds as reservoir hosts but fail to transmit after spillover into mammal hosts such as humans and horses, where they are regularly fatal (Weaver and Barrett, 2004). To investigate this we ran an additional model identifying parasites associated with avian reservoirs, but found no strong evidence that these parasites cause higher mortality (Table 5). It is possible that the positive relationship between virulence and evolutionary isolation breaks down at these larger phylogenetic distances. For example, non-host resistance may be more common following large phylogenetic jumps (van Baarlen et al., 2007). Expanding our framework to include non-mammal hosts may provide additional insight into trade-offs faced by parasites exhibiting extreme phylogenetic generalism.

In evolutionarily isolated hosts, mortality may result from a combination of direct damage caused by parasites, and damage caused by the host’s immune response to infection, which may impose different selective pressures on the evolution of virulence (Graham et al., 2005). Hosts that contribute little to transmission provide one pathway via which transmission can become decoupled from virulence, resulting in parasites experiencing little or no selection to reduce hypervirulence (Antonovics et al., 2013; Woolhouse et al., 2001). This can occur when the majority of transmission is facilitated by a reservoir host, such as has been suggested for foot and mouth disease in southern Africa which uses asymptomatic African buffalo as a reservoir, but causes severe outbreaks after spillover in domestic cattle (Michel and Bengis, 2012). In the example of Rinderpest in Africa, cattle facilitated the sustained transmission of the virus, which caused widespread mortality following spillover into wild ungulates (Barrett and Rossiter, 1999). Many of the domesticated animal diseases we analyze here may represent spillover of infections from wildlife reservoirs, however identifying reservoir species can be challenging and for many parasites included here the reservoir species are unknown.

Virulence may also become decoupled from transmission if parasites infect tissues unrelated to transmission, such as bacterial meningitis infection of the central nervous system (Levin and Bull, 1994), or when parasite stages can persist for long periods of time in the environment (Cressler et al., 2016). While there was no clear relationship between transmission mode and host mortality in our model (Fig 4), parasite identity had an important effect (Table 2, Fig 5, Fig 6), suggesting that other parasite traits modify virulence.

The animal diseases for which we have multiple case-fatality estimates are weighted towards those that have large impacts on international trade. These diseases may more often display high mortality, providing a window into the evolution of virulence that would otherwise be hard to observe. In natural systems, spillover of highly virulent diseases often display stuttering chains of transmission before burning out (Longdon et al., 2014), and thus instances of deadly disease in wildlife may frequently go undocumented (Leggett et al., 2013). High host densities allow parasites to maintain transmission despite causing high mortality (Mennerat et al., 2010), and artificially high densities of domesticated animals may facilitate the maintenance of more deadly diseases, allowing us to better observe their behaviour.

Predicting the outcomes of novel host-parasite interactions presents a major challenge in disease ecology. There is a pressing need to address this challenge given rapid rates of ecosystem transformation which can generate communities never before seen in evolutionary history and promote disease emergence in novel hosts. Proactive approaches to document wildlife hosts (Farrell et al., 2013) may help predict mortality of emerging diseases, and disease burdens may be reduced by implementing effective disease management practices. As a step towards this, we have shown host evolutionary isolation to be a strong predictor of infection-induced mortality in domesticated mammals, and quantify the potential for country-level initiatives to reduce animal death.

## Materials & Methods

Using a global database of infection-induced mortality rates, we employ a Bayesian hierarchical modelling framework to examine the relationship between host specificity and mortality for diseases of domesticated mammals. To separate the importance of our two aspects of host specificity (host evolutionary isolation and host species richness) from other factors that might also influence host mortality, we include co-predictors and hierarchical terms in our model. At the parasite level these include traits for major modes of transmission, plus hierarchical effects of parasite type to account for parasite traits not measured directly. We also include hierarchical effects for host, host taxonomic order, country, and year of reporting. Environmental conditions, which include socio-economic factors such as the ability of local peoples to maintain animal health, effects of ambient temperature on parasite growth rate, or co-infection with other parasites may also influence host mortality. To control for these country-level effects we include per capita Gross Domestic Product (GDP) and latitude per country in addition to modelling variation among countries. The virulence-transmission trade-off suggests that outbreaks resulting in large numbers of infected individuals are unlikely to be associated with high mortality, as premature host death restricts transmission rate, ultimately resulting in lower case numbers for more lethal diseases (Alizon et al., 2009). We therefore also include the number of cases per report as an offset variable. We estimate the effect sizes of these predictors on host mortality with a Bayesian hierarchical binomial-logit model.

### Case-fatality reports

Reports of number of cases and deaths due to infection were taken from published OIE year end reports for the years 2005-2011 (World Organisation for Animal Health (OIE), 2005, 2006, 2007, 2008, 2009, 2010, 2011). Reported by individual countries, these include information per disease-host combination on the number of cases (infected individuals), deaths due to infection, individuals destroyed, and individuals slaughtered. We included only reports of diseases in mammal hosts. We excluded any observations in which host individuals were reported as destroyed or slaughtered as this would interfere with estimates of deaths due to infection. We also excluded the few instances where the reported number of deaths due to infection exceeded the number of reported cases.

### Host and parasite Latin binomials

Reported host codes were assigned a latin binomial based on a combination of geographic location, OIE reports, and classifications defined Clutton-Brock (1999) (Table 1). Reports that included OIE host codes “cer” (cervidae) and “o/c” (sheep or goats) could not be attributed to a single host species and were excluded.

**Table 1:** Conversion table for OIE host codes to latin binomials.

| OIE code | Location | Binomial |
| --- | --- | --- |
| bov | Global | *Bos taurus* |
| buf | Sub-saharan Africa | *Syncerus caffer* |
| buf | North America | *Bison bison* |
| buf | Europe, Latin America, Asia,<br>Caribbean, North Africa | *Bubalus bubalis* |
| can | Global | *Canis lupus* |
| cap | Global | *Capra hircus* |
| cml | Global | *Camelus dromedarius* |
| equ | Global | *Equus caballus* |
| fel | Global | *Felis silvestris* |
| lep | Global | *Oryctolagus cuniculus* |
| ovi | Global | *Ovis aries* |
| sui | Global | *Sus scrofa* |

Reported disease names were assigned a parasite latin binomial based on OIE publications (disease summaries from the OIE Terrestial Manual (World Organisation for Animal Health (OIE), 2012) and OIE technical disease cards). For diseases caused by a particular subspecies or strain, this sub-type was kept in cases where susceptible host species was available (Equine Influenza being largely caused by strain H3N8, and Paratuberculosis caused by *Mycobacterium avium paratuberculosis*). Diseases attributed to multiple species were removed (Atrophic rhinitis of swine, Equine piroplasmosis, Equine rhinopneumonitis, Horse mange, Leishmaniosis, Leptospirosis, Sheep and goat pox, Theileriosis, Trichinellosis, and Trypanosomosis), unless the likely causative species could be identified based on geography and/or reported host species (Bovine babesiosis in Europe caused by *Babesia divergens*, Malignant catarrhal fever in sheep worldwide largely caused by *Macavirus ovine herpesvirus 2*, and Malignant catarrhal fever in African cattle caused by *Macavirus alcelaphine herpesvirus 1*). Diseases caused by prions (Scrapie, Bovine Spongiform Encephalopathy) were excluded.

### Host specificity

The suite of mammalian host species infected by each parasite was taken from the Global Mammal Parasite Database 2.0 (Stephens et al., 2017) and a static version of the Enhanced Infectious Disease Database (EID2) database (Wardeh et al., 2015). Host species for *Influenza A H3N8* and *Mycobacterium avium paratuberculosis* are not included in the static version of the EID2 database, so were instead taken from EID2 online (eid2.liverpool.ac.uk) on June 14^th^ 2017. We also included the host species reported as infected by each parasite in the OIE report data used in the analysis. Host latin binomials were standardized to Wilson and Reeder (2005) using the online version (www.departments.bucknell.edu/biology/resources/msw3) and the Wilson and Reeder 1993-2005 binomial synonym table included in PanTHERIA (Jones et al., 2009). Hosts reported to subspecies were collapsed to the parent binomial, and hosts not reported to species level were removed. *Homo sapiens* were excluded. Host species richness was then calculated as the number of unique host latin binomials associated with each parasite. For each combination of host and parasite reported in the OIE data, mean phylogenetic distances from all known hosts to the infected OIE host was calculated using the Fritz et al. mammal supertree (Fritz et al., 2009) and the R package ape version 3.4 (Paradis et al., 2004).

### Parasite traits

Transmission mode is often listed as a key factor linked to virulence (Alizon et al., 2009; Ewald, 1983; Rigaud et al., 2010; Cressler et al., 2016). Here we include whether a parasite is transmitted by an arthropod vector, is transmitted as a function of reproduction (either vertically transmitted, sexually transmitted, or passed from mother to offspring via ingestion of milk or colostrum), and whether it has a resting stage capable of persisting for long periods of time in the environment (typically months to years). Binary parasite traits coding primary modes of transmission and the use of avian species as reservoir hosts were taken from OIE publications (disease summaries from the OIE Terrestial Manual (World Organisation for Animal Health (OIE), 2012) and OIE technical disease cards), and from Lefèvre et al. (2010). Parasite-level effects were modelled as a function of these covariates plus hierarchical effects of parasite type (virus, bacteria, helminth, etc…), to account for phylogenetic non-independence and capture additional parasite traits not measured directly.

### Country-level covariates

Host mortality is also likely influenced by local environmental conditions. In our data, these may include socio-economic factors such as the ability of local peoples to maintain animal health, effects of ambient temperature on parasite growth rate, or co-infection with other parasites. While the scale of reporting does not allow us to investigate these factors directly, we include two country-level predictors: 1) per capita Gross Domestic Product (GDP) to model economic abilities to reduce host mortality, and 2) latitude as a proxy for temperature and biodiversity gradients that may reflect environmental conditions determining the strength of species interactions (Schemske et al., 2009), in addition to modelling country-level variation. To include country-level covariates from the World Bank World Development Indicators API, we standardized country names to those used in the WDI R package version 2.4 (Arel-Bundock, 2013). For each country we extracted mid-country latitude and per capita in current US dollars (WDI code “NY.GDP.PCAP.CD”) using the WDI package. Countries that did not have reported GPD per capita from the WDI were supplemented with information from the United Nations Data Retreival System (`data.un.org`) so that there was at least one estimate of per capita GDP for the period of 2005-2011. Mean gross domestic product per capita per country was then calculated across all years. We excluded records from countries with no iso3 code or for which no latitude was reported.

### Model

Using a hierarchical Bayesian binomial-logit model, we model deaths (*deaths*_*i*_) as following a binomial distribution determined by sample size per observation (*cases*_*i*_) and a probability parameter *p*_*i*_. The higher-level structure of the model is as follows:

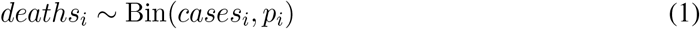

Where *p*_*i*_ is modeled with *β*_0_ as the grand mean plus the effects of mean phylogenetic distance from all known hosts to the species infected (*EvoIso*_*i*_), the number of cases per observation (*cases*_*i*_), and partially-pooled hierarchical effects for parasites (*μ*_*para*_), hosts (*μ*_*host*_), countries (*μ*_*country*_), and years (*μ*_*year*_):

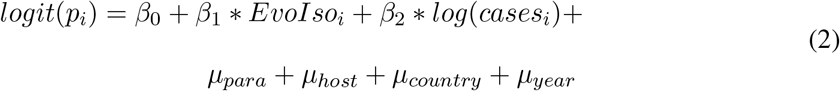

Parasite level effects, *μ*_*para*_, are defined by a normal distribution as follows:

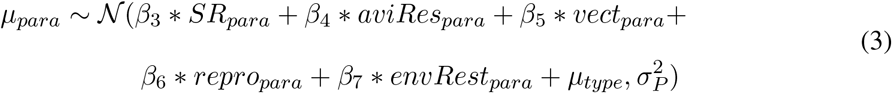

Where the difference from the grand mean (*β*_0_) for each parasite (*para*) is determined by host species richness (*SR*_*para*_), transmission modes (*aviRes*_*para*_, *repro*_*para*_, *envRes*_*para*_), and a hierarchical effect of the parasite type (*μ*_*type*_), and variance parameter 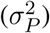.

Parasite taxonomic type (i.e. virus, bacteria, helminth, etc…), *μ*_*type*_, is modelled following a normal distribution with mean of zero and variance parameter 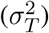 as follows:

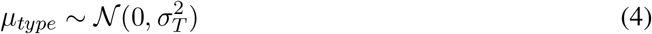

Host level effects, *μ*_*host*_, are modelled following a normal distribution with mean determined by a hierarchical effect of the host taxonomic order (*μ*_*order*_) and variance parameter 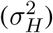 as follows:

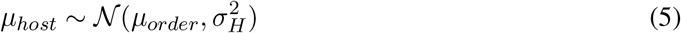

Host taxonomic order level effects, *μ*_*order*_, are modelled following a normal distribution with mean of zero and variance parameter 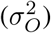 as follows:

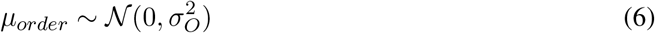

Country level effects, *μ*_*country*_, are modelled following a normal distribution with mean determined by gross domestic product per capita (*GDP*_*c*_) and latitude (*latitude*_*c*_), and variance parameter 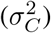 as follows:

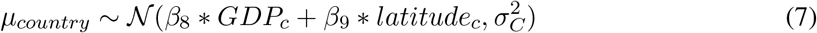

Year level effects, *μ*_*year*_, are modelled following a normal distribution with mean of zero and variance parameter 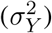 as follows:

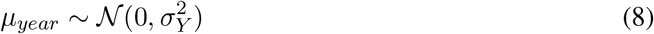

### Priors & Data transformations

Following the recommendations of Gelman et al. (2008), continuous predictors were normalized to mean of zero and standard deviation of 0.5. Estimated parameters were modelled using weakly informative priors as recommended by Ghosh et al. (2015) and the Stan development team:

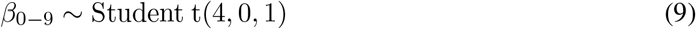

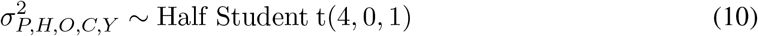

### Sampling and Convergence Diagnostics

Models were fit in Stan (The Stan Development Team, 2017; Carpenter et al., 2017) via R 3.2.3 (R Core Team, 2015) with rstan version 2.14.2 (Stan Development Team, 2017a) using 4 chains with 30,000 iterations per chain. The first 15,000 iterations per chain were used for warm-up and discarded. The remaining posterior was thinned to retain every 10th iteration, resulting in a total of 6,000 posterior draws. Convergence was diagnosed by observation of Rhat values equal to 1 (Table 2) and explored with shinystan version 2.4.0 (Stan Development Team, 2017b). Posterior predictive checks were performed to ensure model validity and fit to the data. The main model was also fit with simulated data to ensure the model performs as expected and is able to recover simulated parameters.

## Acknowledgments

We thank Vanessa Ezenwa, Charlie Nunn, Elizabeth Wolkovich, Ria Ghai, Jan Gogarten, and Carl Boodman for helpful feedback on the manuscript. Special thanks are due to Elizabeth Wolkovich, Margaret Kosmala, the Harvard Stanleyi group, Will Pearse, and Bob Carpenter for feedback on the model, and to Debarun Gupta & Madeleine McGreer for help with data entry. MJF was supported by a Vanier NSERC CGS, the CIHR Systems Biology Training Program, the Quebec Centre for Biodiversity Science, and the McGill Biology Department.

## Supplementary Information

### Main model

**Table 2:** Summary of model output for continuous predictors including posterior means, posterior standard deviations, 2.5%, 25%, 50%, 75% and 97.5% quantiles, the effective sample size (n_eff), and the potential scale reduction statistic (Rhat).

| Level | Parameter | mean | sd | 2.5% | 25% | 50% | 75% | 97.5% | n_eff | Rhat |
| --- | --- | --- | --- | --- | --- | --- | --- | --- | --- | --- |
|  | Intercept | -0.57 | 0.74 | -2.06 | -1.03 | -0.57 | -0.08 | 0.87 | 1338 | 1.00 |
|  | log (Cases) | -1.33 | 0.01 | -1.35 | -1.34 | -1.33 | -1.32 | -1.30 | 6000 | 1.00 |
|  | Evolutionary Isolation | 1.69 | 0.12 | 1.45 | 1.61 | 1.70 | 1.78 | 1.93 | 6000 | 1.00 |
| 4*Parasite | Host Species Richness | 0.90 | 0.71 | -0.42 | 0.41 | 0.88 | 1.37 | 2.36 | 3335 | 1.00 |
|  | Vectored | -0.27 | 0.65 | -1.60 | -0.69 | -0.26 | 0.15 | 1.00 | 3631 | 1.00 |
|  | Reproduction | 0.12 | 0.66 | -1.19 | -0.32 | 0.13 | 0.56 | 1.43 | 4317 | 1.00 |
|  | Environmental Resting Stage | 0.22 | 0.96 | -1.65 | -0.40 | 0.20 | 0.83 | 2.19 | 6000 | 1.00 |
| 2*Country | Latitude | -0.21 | 0.42 | -1.04 | -0.49 | -0.21 | 0.08 | 0.61 | 3796 | 1.00 |
|  | GDP per capita | -0.56 | 0.37 | -1.29 | -0.81 | -0.56 | -0.31 | 0.18 | 6000 | 1.00 |

**Figure 5:**
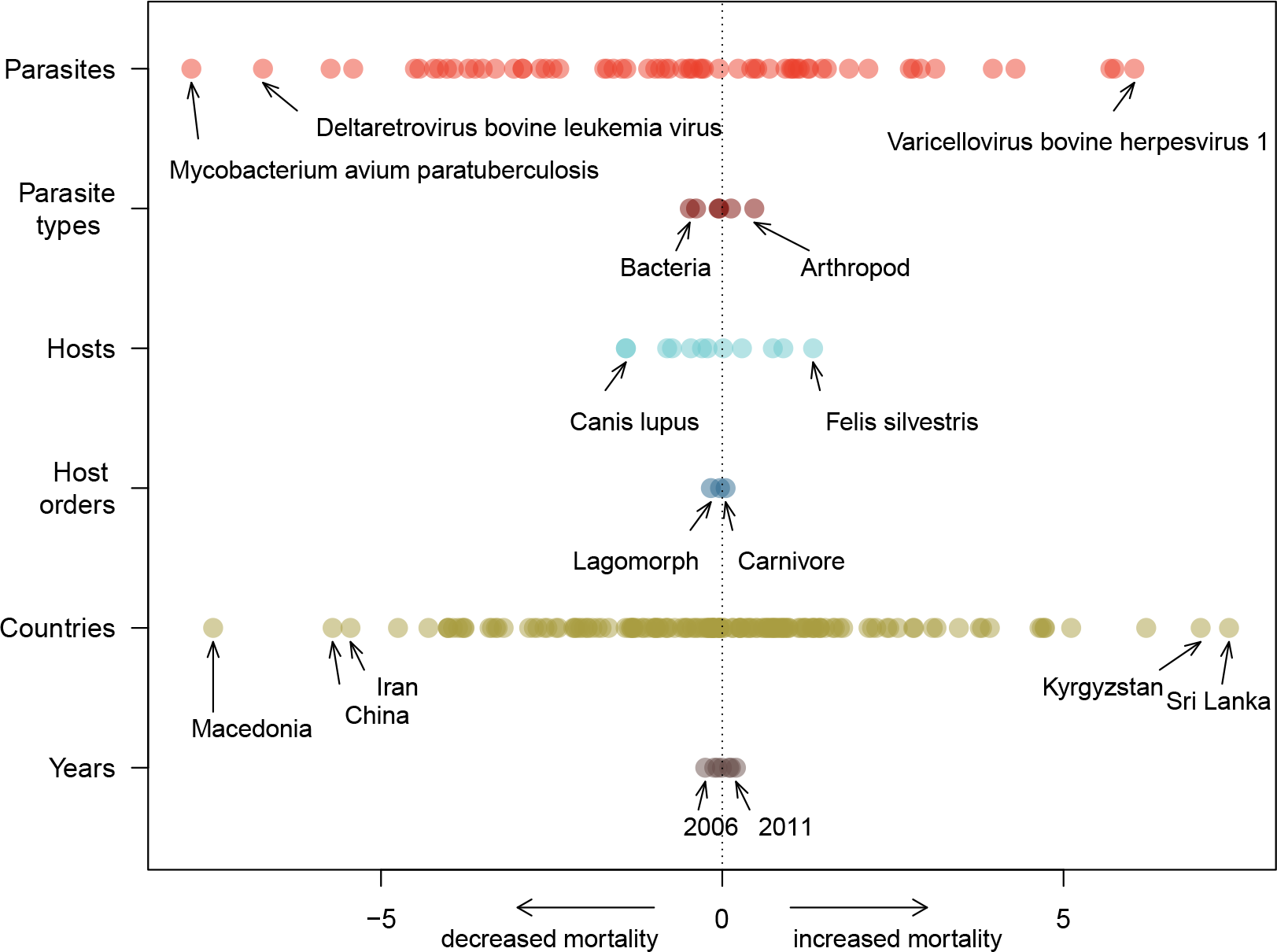
Mean estimated effects for individual hierarchical terms (parasites, parasite types, hosts, host orders, countries, and years). Plotted estimates have been set to 50% transparency to visualize overlapping points, and extreme estimates in each group have been identified.

**Figure 6:**
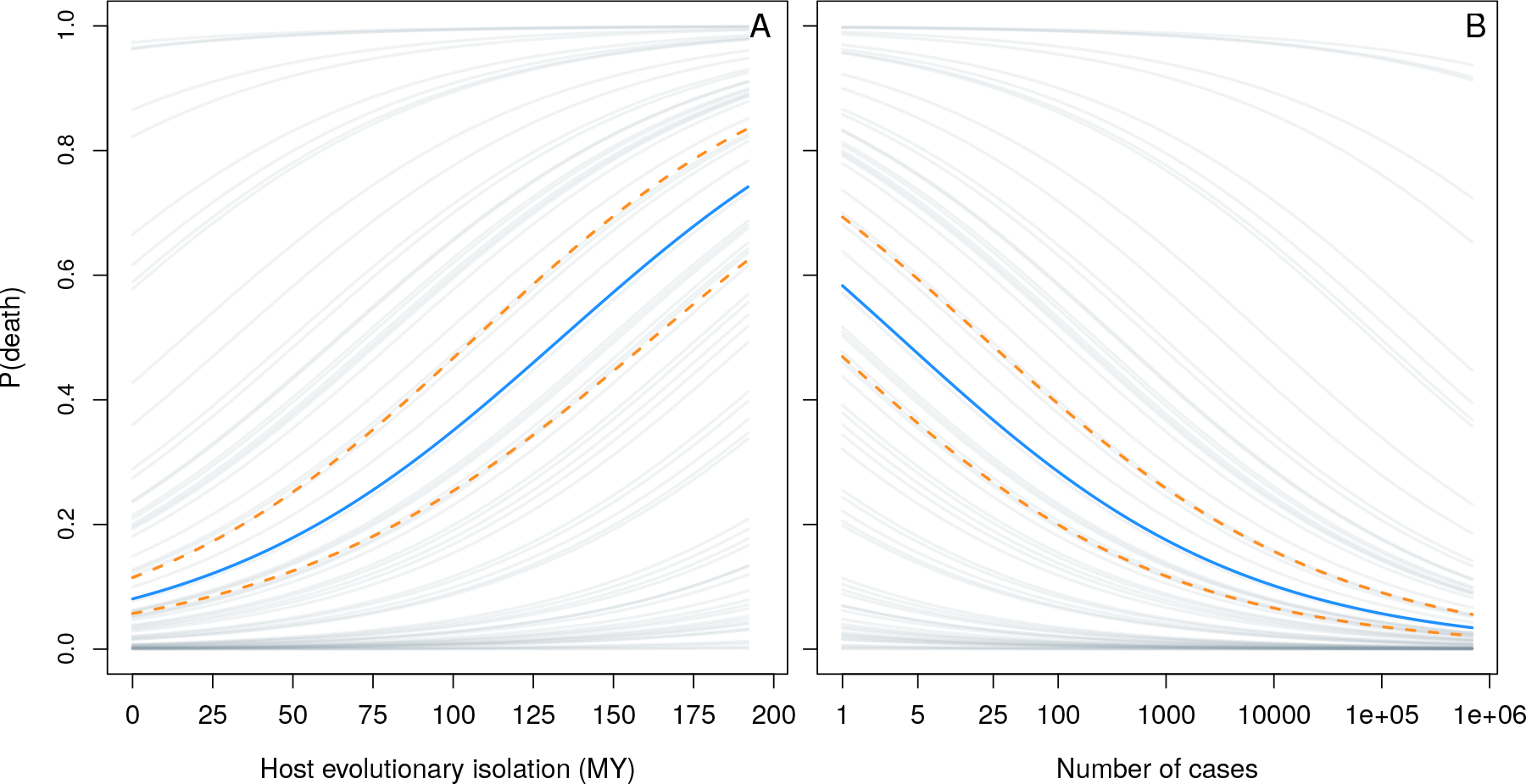
Posterior predictions of the probability of death as a function of A) host evolutionary isolation (in millions of years), and B) the number of cases. Solid blue lines represent the mean logistic curve, dashed yellow lines represent the upper and lower bounds of the 50% credible interval. Grey lines depict equivalent mean curves offset by the posterior mean effects for each parasite.

### Sensitivity Analyses and Alternative Models

#### Excluding single-host parasites

As selective pressures driving virulence evolution are likely to differ among single and multi-host parasites, the main model fit again after removing single-host parasites from the data.

**Table 3:**
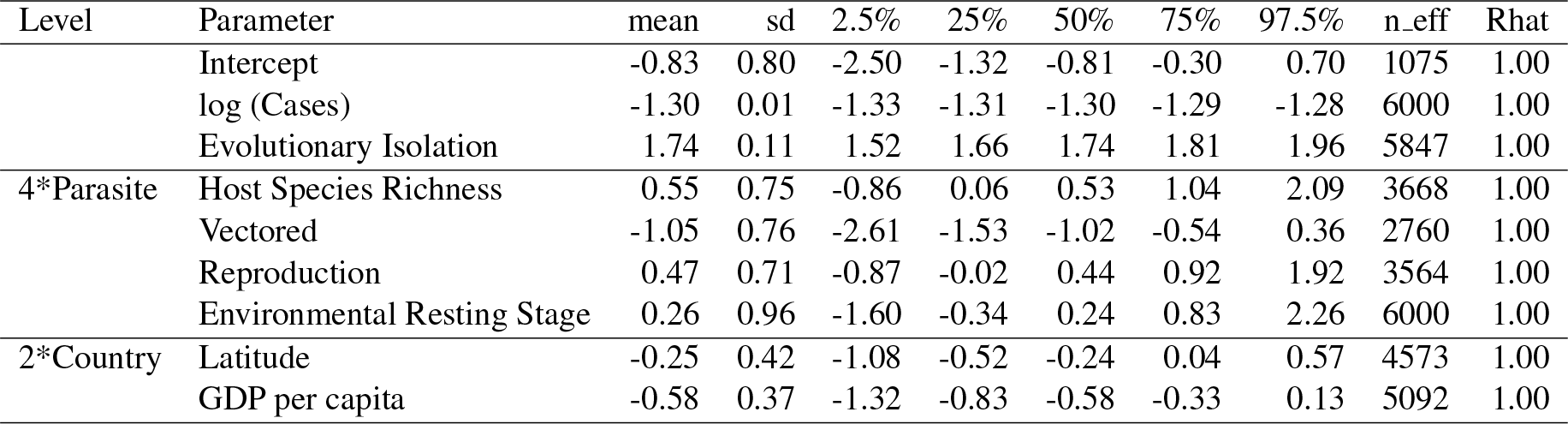
Summary of main model excluding single-host parasites for continuous and binary predictors including posterior means, posterior standard deviations, 2.5%, 25%, 50%, 75% and 97.5% quantiles, the effective sample size (n_eff), and the potential scale reduction statistic (Rhat).

#### Host taxonomic diversity

Due to incomplete sampling, the host species reported in the GMPD and EID2 databases are unlikely to include the complete set of susceptible hosts for each parasite. As a sensitivity analysis, host species richness (*SR*_*p*_) was replaced by a measure of taxonomic diversity using data reported Lefèvre et al. (2010) and the OIE documentation. Host taxonomic diversity varies from 1-6 corresponding to whether parasites infect hosts belonging to a single species (1), genus (2), family (3), order (4), class (5), or multiple classes (6). Just as with host species richness, the ability to infect humans was not included in estimates of taxonomic diversity.

**Table 4:**
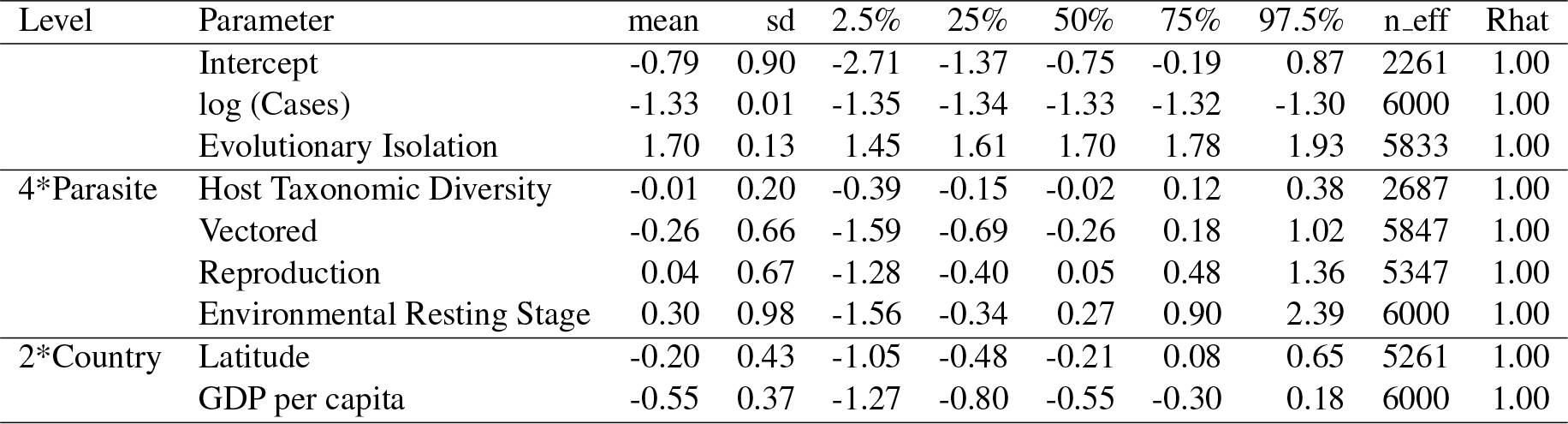
Summary of model with host taxonomic diversity for continuous and binary predictors including posterior means, posterior standard deviations, 2.5%, 25%, 50%, 75% and 97.5% quantiles, the effective sample size (n eff), and the potential scale reduction statistic (Rhat).

#### Parasites with avian reservoirs

As an extension of our main model, we include whether or not a parasite uses an avian reservoir (Eastern equine encephalitis, Western equine encephalitis, Venezuelan equine encephalitis, Fowlpox, Newcastle Disease, West Nile Virus, *Pasturella multocida*), as we hypothesize that this might cor-relate with whether domesticated mammals represent dead-end hosts from which the parasite is not transmitted further, such as is the case for West Nile Virus and other encephalitic viruses that spillover from birds to horses (Weaver and Barrett, 2004). The use of avian species as reservoir hosts were taken from OIE publications (disease summaries from the OIE Terrestial Manual (World Organisation for Animal Health (OIE), 2012) and OIE technical disease cards), and from Lefèvre et al. (2010), and coded as a binary predictor.

**Table 5:** Summary of model including indicator for avian reservoir for continuous and binary predictors including posterior means, posterior standard deviations, 2.5%, 25%, 50%, 75% and 97.5% quantiles, the effective sample size (n eff), and the potential scale reduction statistic (Rhat).

| Level | Parameter | mean | sd | 2.5% | 25% | 50% | 75% | 97.5% | n_eff | Rhat |
| --- | --- | --- | --- | --- | --- | --- | --- | --- | --- | --- |
|  | Intercept | -0.59 | 0.72 | -2.04 | -1.07 | -0.59 | -0.11 | 0.80 | 1568 | 1.00 |
|  | log (Cases) | -1.33 | 0.01 | -1.35 | -1.33 | -1.33 | -1.32 | -1.30 | 5980 | 1.00 |
|  | Evolutionary Isolation | 1.69 | 0.12 | 1.46 | 1.61 | 1.69 | 1.78 | 1.93 | 5935 | 1.00 |
| 4*Parasite | Host Species Richness | 0.90 | 0.73 | -0.45 | 0.41 | 0.87 | 1.36 | 2.42 | 3574 | 1.00 |
|  | Vectored | 0.28 | 0.65 | -1.60 | -0.70 | -0.26 | 0.16 | 1.01 | 4318 | 1.00 |
|  | Reproduction | 0.14 | 0.65 | -1.14 | -0.30 | 0.13 | 0.58 | 1.42 | 5209 | 1.00 |
|  | Environmental Resting Stage | 0.23 | 0.97 | -1.61 | -0.39 | 0.20 | 0.82 | 2.29 | 5934 | 1.00 |
|  | Avian Reservoir | 0.18 | 0.83 | -1.47 | -0.36 | 0.17 | 0.68 | 1.83 | 5474 | 1.00 |
| 2*Country | Latitude | -0.20 | 0.42 | -1.05 | -0.48 | -0.20 | 0.09 | 0.62 | 3896 | 1.00 |
|  | GDP per capita | -0.55 | 0.37 | -1.29 | -0.80 | -0.55 | -0.31 | 0.15 | 5942 | 1.00 |

**Figure 7:**
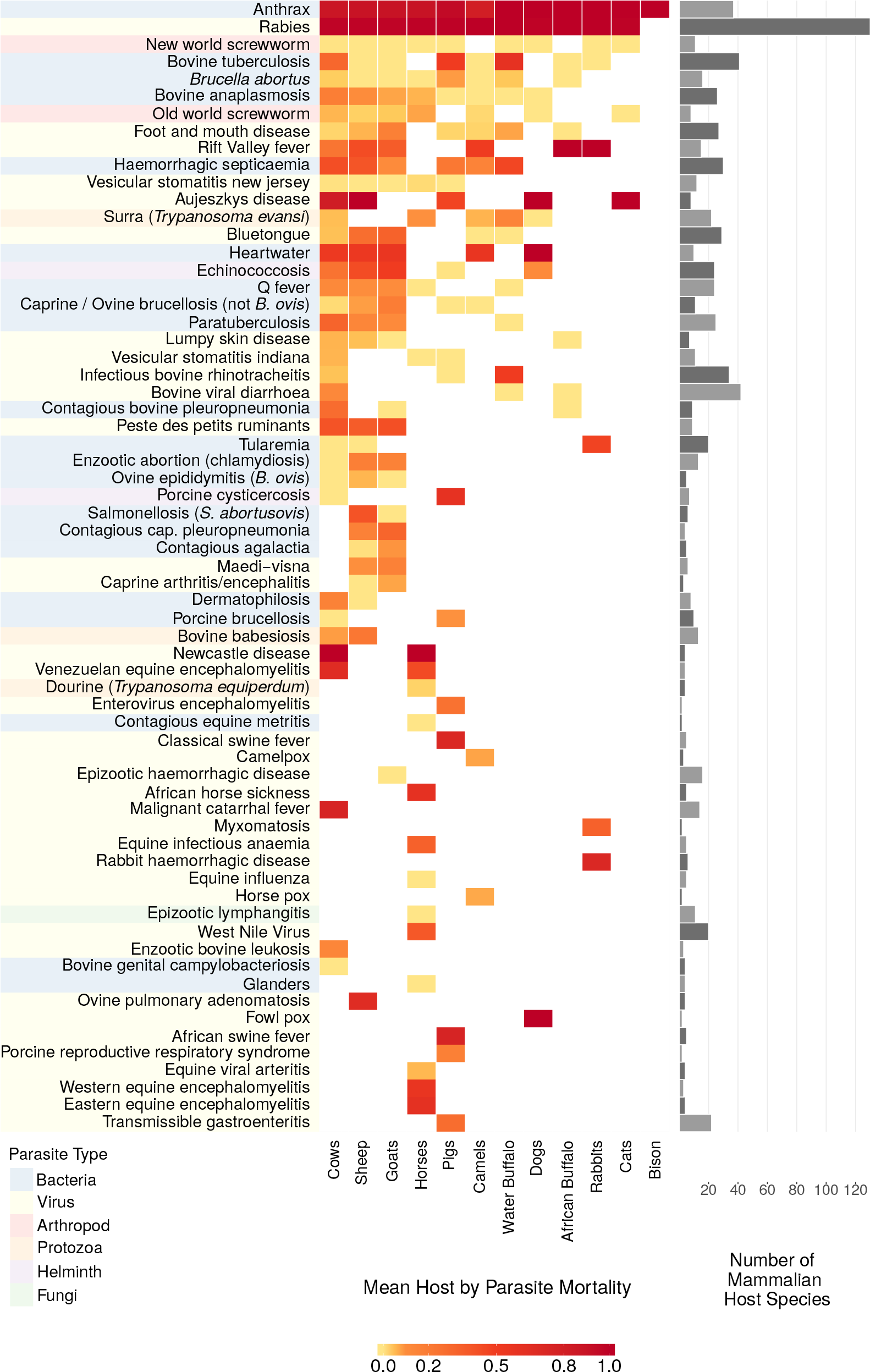
Version of main document Fig. 2 including parasite common names. Parasite names are colour coded by parasite type.

